# Uncovering Additional Predictors of Urothelial Carcinoma from Voided Urothelial Cell Clusters Through a Deep Learning Based Image Preprocessing Technique

**DOI:** 10.1101/2022.04.30.490136

**Authors:** Joshua J. Levy, Xiaoying Liu, Jonathan D. Marotti, Darcy A. Kerr, Edward J. Gutmann, Ryan E. Glass, Caroline P. Dodge, Arief A. Suriawinata, Louis J. Vaickus

## Abstract

Urine cytology is commonly used as a screening test for high grade urothelial carcinoma for patients with risk factors or hematuria and is an essential step in longitudinal monitoring of patients with a prior bladder cancer history. However, the semi-subjective nature of current reporting systems for urine cytology (e.g., The Paris System) can hamper reproducibility. For instance, the incorporation of urothelial cell clusters into the classification schema is still an item of debate and perplexity amongst expert cytopathologists, as several previous works have disputed their diagnostic relevance. Recently, several machine learning and morphometric algorithms have been proposed to provide quantitative descriptors of urine cytology specimens in an effort to reduce subjectivity and include automated assessments of cell clusters. However, it remains unclear how these computer algorithms interpret/analyze cell clusters. In this work, we have developed an automated preprocessing tool for urothelial cell cluster assessment that divides urothelial cell clusters into meaningful components for downstream assessment (i.e., population-based studies, workflow automation). Results indicate that cell cluster atypia (i.e., defined by whether the cell cluster harbored multiple atypical cells, thresholded by a minimum number of cells), cell border overlap and smoothness, and total number of clusters are important markers of specimen atypia when considering assessment of urothelial cell clusters. Markers established through techniques to separate cell clusters may have wider applicability for the design and implementation of machine learning approaches for urine cytology assessment.

## Introduction

Urine cytology (UC) specimens are essential for bladder cancer detection and screening yet are challenging to assess given the complexity of the specimen. Screening is often accomplished through application of The Paris System (TPS) criteria, which assigns four main ordered categories (negative, atypical urothelial cells, suspicious for high grade urothelial carcinoma, positive for high-grade urothelial carcinoma, HGUC) based on the following criteria for a positive diagnosis: 1) at least 5 malignant urothelial cells, 2) a nuclear-to-cytoplasm (NC) ratio at or above 0.7, 3) nuclear hyperchromasia, 4) markedly irregular nuclear membrane, and 5) coarse/clumped chromatin ^1,2^. Specimens with definitive diagnoses (negative, positive) are often easier to assess than atypical (hedged against negative diagnosis) and suspicious (hedged against positive diagnosis; less than five malignant cells needed) specimens. Predictably, these two indeterminate diagnoses are hindered by poor interobserver variability as compared to negative or positive diagnoses ^3–5^.

TPS does not explicitly establish urothelial cell clusters as separate assessment criteria for the final cytologic diagnosis. Instead, clusters are judged by their constituent cells, where cytomorphological assessments for each cell in the cluster must satisfy atypical criteria and combined with the individual cell assessments. However, the significance of cell clusters for urothelial cancer has not been fully elucidated. Whether and how urothelial clusters are assessed during bladder cancer screening potentially impacts diagnostic reproducibility. Several studies have found no association between number and type of cell cluster and urothelial carcinoma, whereas others have demonstrated a statistically significant positive association between the number of clusters and urothelial carcinoma ^2,6–10^. One such study established three architectural types of urothelial cell clusters: 1) non-overlapping cells, 2) overlapping cells, densely packed, lacking distinguishable cell borders (dense regions), and 3) overlapping cells with distinguishable cell borders ^6^. While the study authors were able to identify that the number of clusters, irrespective of cluster type, were associated with specimen atypia, presence of urothelial carcinoma was not associated with any cluster type alone. As another example, tissue fragments and papillary clusters with a fibrovascular core as an indication for urothelial carcinoma (i.e., low-grade carcinoma) yet are rare. Presence and type of cell clusters and tissue fragments in urine specimens may also be an artifact of specimen preparation (e.g., ThinPrep® associated with higher presence of clusters), previous reports of tissue biopsy, urothelial stones, etc., all of which potentially impact exfoliation of cell clusters. Specimens containing cell clusters are typically classified as atypical, favor reactive or low grade urothelial neoplasm diagnosis, even without the fibrovascular core ^2^. Previous studies have deemphasized the diagnostic utility of urothelial clusters in favor of individual cell analysis ^8,11,12^. However, prevalent diagnostic criteria have been established almost exclusively based on inspection of voided specimens, while significant differences have been documented between types of specimen preparation (e.g., cystoscopic, voided, upper tract, instrumented, neobladder, etc.).

Several computational, image analysis and machine learning methods have been developed to quantitatively assess urine cytology specimens to generate an automated summary ^13–21^. In brief, these methods can parse digitized representations of cytology slides (Whole Slide Images; WSI) into their constituent cellular components. Parsed cells are independently assessed using morphometric and machine learning approaches to quantify features of atypia (e.g., NC ratio, hyperchromasia, clumped chromatin, etc.). Features are then tabulated across all cells in the slide to measure the overall atypia burden of the specimen in order to predict the presence of high-grade urothelial carcinoma. Automation tools for urine cytology could provide both rapid and objective screening and have the potential to improve diagnostic accuracy and disease management by assessing every cell in a given specimen, a prescreening task normally assigned to cytotechnicians prior to final assessment from the cytopathologist^22^. Cytotechnologists, although ostensibly expected to screen every cell on a given slide, realistically do not perform such an exhaustive assessment. As compared to cytopathologists, they perform a more regimented screening, combining single cell assessment and gestalt impressions over the entire slide area.

While existing image assessment techniques have either uncovered new or recapitulated previously known specimen malignancy predictors, the handling of clusters is still inadequately addressed. For instance, a previous machine learning technique used density-based clustering methods to identify cell clusters, which were then validated using a neural network. However, the sensitivity and specificity of the clustering method for identifying cell clusters was not validated in the paper and information was not explicitly provided on how clusters factored into the final assessment other than to report the number of clusters in the specimen ^13^. In addition, associations between cluster scores and outcomes were not made available. Another technique used nucleus-centric watershed-based methods to segment urothelial cells for the calculation of a cluster-specific NC ratio ^14^. Such techniques assume cell borders do not overlap and may provide imprecise atypia estimates. Furthermore, the performance of the water shedding technique to delineate urothelial cells from other cell types in the cluster was not discussed. Nonetheless, these methods present a significant advancement from previous modes of assessment.

In this study, we detail the development of an Artificial Intelligence (AI) tool for urothelial cell cluster analysis that can accurately localize urothelial cells, overlapping cell boundaries, dense regions of significant overlap and identify visual markers of urothelial atypia. By breaking clusters into their constituent architectural components ^6^ and isolating cells with overlapping cell borders, this preprocessing tool can facilitate downstream association studies and development of predictive algorithms that explicitly incorporate cluster architecture and assess the cytomorphology of overlapping cells for atypia. As a demonstration of this approach, we use our tool to associate the quantitative cluster-level features with high-grade urothelial carcinoma. The goal of this work was to build more precise quantitative descriptors of urine clusters. In the future, we plan to use the cluster tool as a preprocessing mechanism for an automated workflow that enables rapid screening of all types of cytology specimens.

## Methods

### Data Collection and Image Scanning

We collected 1,277 urine specimens across 141 patients from 2008 to 2019 at our institution, Dartmouth-Hitchcock Medical Center (DHMC). Specimens were prepared through ThinPrep® and Pap-staining prior to microscopic examination. Urine slides were scanned using a Leica Aperio-AT2 scanner at 40x resolution and stored as full-resolution SVS files representing WSI.

### Annotation of Cell Clusters for Training, Validation and Internal Test Set

Candidate cell clusters were separated from background in each WSI through a connected components analysis that assigns each cell/cluster a separate identifier. For each cell/cluster, remaining background was masked out and images of each candidate cluster were stored as PNG/TIFF files (**Figure 1A**). Candidate cell clusters were subsequently filtered based on cluster size and number of predicted nuclei using an updated version of a previously developed nuclei segmentation tool configured for NC ratio estimation ^14^. Two cytopathologists (LJV and XL) were presented with 800 candidate cluster images for annotation that were not utilized for held-out evaluation. Of the 800 candidate clusters, 633 were confirmed urothelial clusters. The cytopathologists annotated each of these cell clusters by outlining all cell boundaries, where possible, and individual cells were assigned to the following classes: 1) squamous (red), 2) inflammatory (green), 3) negative urothelial cell (purple), 4) atypical urothelial cell (blue), and 5) dense regions of overlapping yet indistinguishable cell borders (orange) which were circled as a group by the cytopathologist (**Figure 1B**). Of the images used to develop the urothelial cell cluster algorithm, 474 clusters were partitioned to the training/validation dataset and 159 clusters served as the internal test set, corresponding to a total 8,123 cells annotated (**Table 1**). Both cytopathologists were given Microsoft Surface tablets and a touch pen to annotate cells, by circling the cell borders and tagging with the relevant cell type / architecture as aforementioned. Note that cell boundaries were annotated here, which is entirely different from the task of annotating cell nuclei. Annotation of cell borders was done using ASAP software (https://computationalpathologygroup.github.io/ASAP/).

**Table 1:** Break down of training/validation and internal test set clusters; by number of clusters and number of annotations for each cell type across clusters

| Cell Type Annotated | Training/Validation Sets | Internal Test Set |
| --- | --- | --- |
| Negative Urothelial Cell | 4050 | 1546 |
| Atypical Urothelial Cell | 568 | 144 |
| **Dense Architecture** | 203 | 83 |
| **Squamous Cells** | 724 | 231 |
| **Inflammatory Cells** | 459 | 115 |
| **Total Cells Annotated** | 6004 | 2119 |
| **Total Clusters** | 474 | 159 |

**Figure 1:**
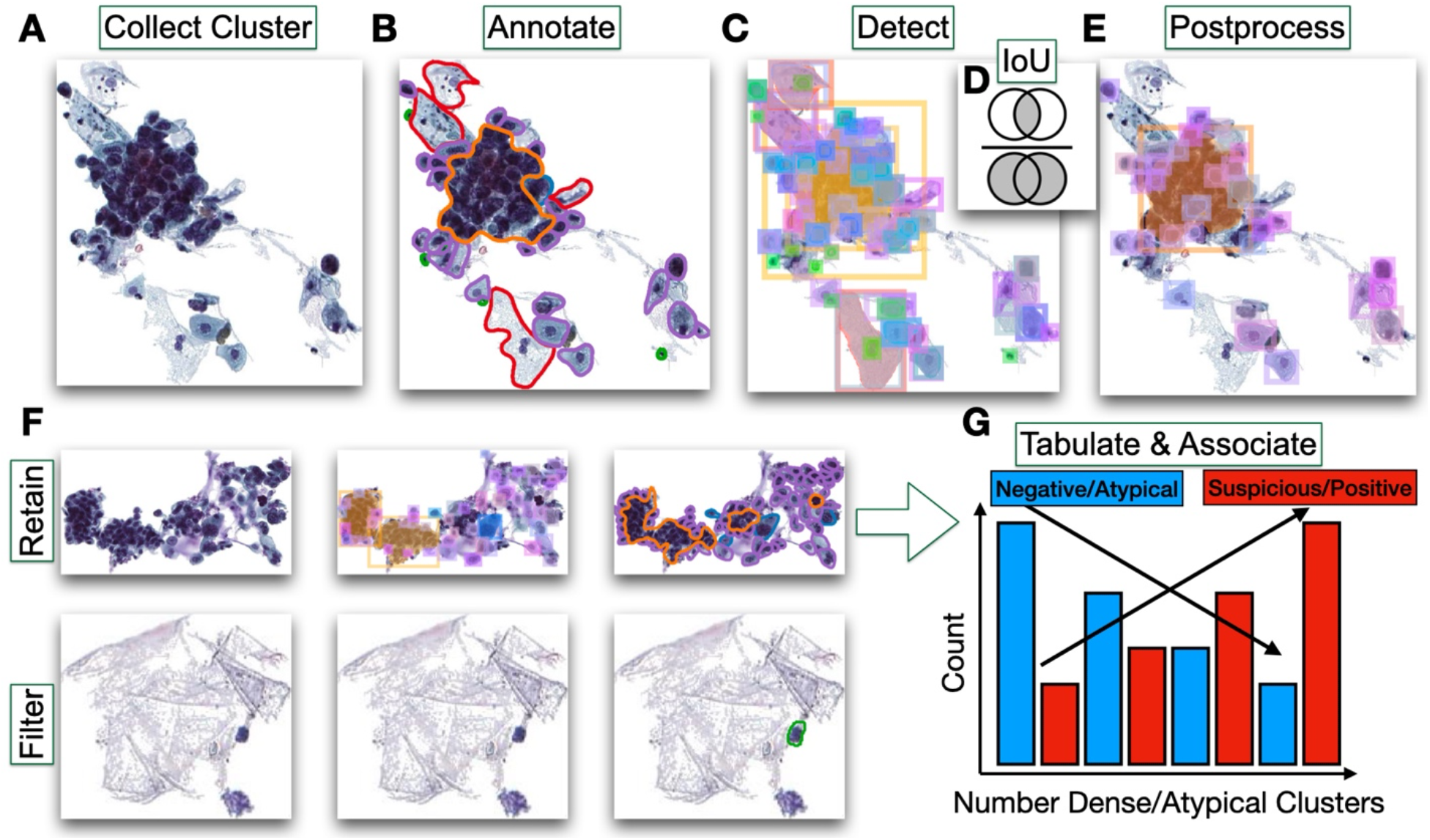
Methods Overview: **A)** Training, validation and internal test set clusters are collected and **B)** annotated for squamous (red), inflammatory (green), negative urothelial cell (purple), atypical urothelial cell (blue), and dense regions (orange); **C)** Cell border detection model identifies candidate cells cytoplasmic borders within cluster; **D)** significantly overlapping cells types, as defined by their intersection over union (IoU), are **E)** filtered by their objectiveness score (e.g., squamous cell predicted in same area as urothelial cell but with higher score) and remaining predicted squamous and inflammatory cells are additionally removed to reveal negative and atypical urothelial cells and dense regions of urothelial cells; **F)** urothelial clusters are called if the number of remaining elements exceed the *minimum cell number*; **G)** number of clusters (urothelial, atypical, dense) are totaled per specimen and totals are then tabulated across the specimens to reveal associations with UC atypia

### Object Detection Model Training and Postprocessing for Training/Validation Clusters

For each candidate cell cluster, we aimed to localize individual cells (squamous, inflammatory, negative/atypical urothelial cells) and cluster architectures (dense regions of overlapping urothelial cells without smooth cell borders), while locating the cell borders, even if overlapping. We were able to accomplish this aim through training of an object detection neural network ^23–26^. In brief, this neural network simultaneously detects multiple objects or *instances* in the image by proposing and filtering regions of interest (i.e., boxes), providing an “objectness” score between 0-1 that ascribes confidence in its prediction, then “tags” each object dynamically with the appropriate label, and finally outlines the object borders using a segmentation neural network, that predicts within the object on a pixel-by-pixel basis the object’s precise boundary (**Figure 1C**). In contrast, water shedding and DBSCAN are unable to perform these tasks and lack the precision required to locate cells/dense architectures which may or may not have overlapping boundaries. Our object detection network was trained on the training/validation set using the Detectron2 framework ^27^. The model was trained for 1000 epochs using NVIDIA V100 GPUs. Training images were augmented (e.g., randomly flipped, rotated, resized, color jitter) during training to improve the model fitness.

After model training, predicted clusters underwent post-processing to remove potentially spurious predictions. Briefly, regions of interest for separate objects may overlap, which is helpful for identifying urothelial cells with overlapping boundaries. However, cell calls with significant overlap, more so than expected with overlapping cytoplasmic boundaries, were filtered using non-maximum suppression (NMS) ^28^, which selects the cell type with the greatest “objectness score” where two objects overlapped (**Figure 1D-E**). Detected squamous cells and inflammatory cells were removed during this filtering process to focus the assessment on urothelial cell atypia and architecture (**Figure 1E**).

### Evaluation of Internal Test Set

Validation of the internal test set clusters were accomplished by calculating concordance statistics between the cytopathologists’ cellular annotations and the model’s predictions. This was accomplished in several ways.

First, we calculated cell-level statistics for the internal test set of clusters: *how well did individually localized cells and dense regions align with the annotations?* To this end, we calculated the average percentage of urothelial cells per cluster that accurately aligned to ground truth urothelial cell annotations, as defined by the *intersection over union* (IoU) score (**Figure 1D**) to associate each ground truth annotation with the prediction with the greatest overlap. Then, for each cluster, we merged all detected dense regions and calculated the average IoU between the predicted and ground truth dense regions, weighted by the size of each dense region, as a measure of detection accuracy. Finally, we assessed the sensitivity and specificity of the approach for instances where the detection model may have conflated non-overlapping and overlapping urothelial cells with dense overlapping regions through cross tabulating predicted urothelial cells and dense regions with their associated ground truth annotations.

Aside from ensuring that the cell-level predictions were accurately calibrated, we also sought to assess cluster level measures (e.g., *does the cluster exhibit cytological and/or architectural atypia*?) for the internal test set. We wanted to evaluate whether these metrics were associated with urothelial carcinoma assignments across the held-out test WSI. Assuming our cell border detection tool was well attuned, our tool could estimate associations that are not subject to rater subjectivity. First, we calculated the number of urothelial cells per cluster, predicted and actual. We then calculated the spearman’s correlation coefficient to measure agreement between the number of predicted and true urothelial cells per cluster. We additionally measured the ability to detect a urothelial cluster (as codified by the ability to accurately detect at least 3 urothelial cells) through calculation of the C-statistic. Similar C-statistics were calculated for the ability to detect whether the cluster was negative (i.e., at least 3 negative urothelial cells) and atypical (i.e., at least 3 atypical urothelial cells). Finally, concordance between predicted and ground truth dense regions were established through calculation of the spearman’s correlation coefficient for area estimates of the dense regions and calculation of the C-statistic for the ability to detect whether the cluster contained a dense region.

We calculated 95% confidence intervals for all cell-level and cluster-level concordance statistics using 1000-sample non-parametric bootstrapping.

### Curation and Automated Cytological Assessment of Clusters in the Held-Out Test Set

Staff cytopathologists assessed the original urine cytology glass slides for evidence of urothelial carcinoma. Olympus microscopes were utilized for pathologic examination. Cytology specimens were assigned primary diagnoses at the time of collection by one of 10 cytopathologists, and secondarily from a reassessment while utilizing The Paris System (TPS) criteria from one of 5 cytopathologists (note: no two cytopathologists assessed the same slide). TPS was officially adopted by DHMC in 2018, so TPS criteria was applied during reexamination of the specimens. In order to eliminate confounding from specimen preparation, only voided urine specimens were considered. Slides that were nondiagnostic were removed from the external assessment, as were slides that contained scanning artifacts and other artifacts that could impact manual and digital assessment (e.g., abundant blood, neobladder, etc.). Specimens corresponding to WSI with an excess of 5 million objects (much of them corresponding to debris) during WSI preprocessing were removed from the analysis. A representative sample of WSI were selected from this dataset. We assessed a total of 430 WSI spanning across 105 patients (**Table 2**). All image processing techniques were implemented in Python v3.7 and large-scale image processing was accomplished in parallel using high-throughput job submission via the Dartmouth College Discovery Research Computing Cluster.

**Table 2:** Breakdown of diagnostic assignments in held-out test set and patient demographics.

|  |  |  |  |  |  |
| --- | --- | --- | --- | --- | --- |
| **History (%)** | **N (%)** | **Type (%)** | **N (%)** | **Age (Mean (SD))** | 73.64 (9.81) |
| **Bladder Cancer** | 363 (84.4) | **Negative** | 232 (54.0) | **Number Sex = M (%)** | 82 (78.1) |
| **Hematuria** | 60 (14.0) | **Atypical** | 131 (30.5) | **Number of Patients** | 105 |
| **Other** | 7 (1.6) | **Suspicious** | 43 (10.0) |  |  |
| **Total Number Specimens** | 430 | **Positive** | 24 (5.6) |  |  |

For each WSI, in contrast to previous works, we included urothelial cell clusters whose predicted number of urothelial cells surpassed a specific threshold (denoted as *minimum cell number*; a proxy for the overall sizes of urothelial clusters considered in the analysis) (**Figure 1F**). Clusters were labeled as *atypical* if they harbored a minimum number of atypical cells (i.e., at least 10% of the *minimum cell number*). Clusters were labeled as *dense* if they contained a dense region of overlapping urothelial cells without a definitive cell border. For a given *minimum cell number*, we tabulated the total number of clusters, atypical clusters, and dense clusters for a given specimen to form slide-level cluster measures. Based on the cytopathologist’s rating, we also dichotomized whether the specimen was assessed to be high risk (suspicious or positive), which carry certain disease management implications (e.g., ordering of cystoscopy or biopsy for histological diagnosis), as opposed to low risk (atypical and negative), which may only necessitate longitudinal follow-up (**Figure 1G**).

### Associations between Cluster Metrics and Cytological Atypia on the Held-Out Test Set

We implemented several Bayesian hierarchical models, fit using Markov Chain Monte Carlo methods ^29^, to associate malignancy (whether specimen was deemed suspicious or positive) with the number of clusters, atypical clusters, and dense clusters (**see Appendix, section “Modeling Number and Type of Urothelial Clusters”**). We did not control for other potential confounders other than what was done through restriction of the final WSI set (detailed above). These models reported incidence rate ratios (*IRR*) and odds ratios (*OR*) and corresponding credible intervals (CI; similar to confidence interval) to describe the association between specimen atypia (suspicious or positive) and number/type of cell cluster. A CI greater than one indicates a positive relationship, whereas a CI less than one indicates a negative relationship. These CI were complemented by calculation of a “p-value”-like measure using the probability of direction, *pd (p* ≈ 2 * (1 − *pd*)) for reporting the existence of a positive/negative association. Model fitting was accomplished using the R v4.1 statistical language via the *brms* package ^30^. *IRR/OR* were reported for different *minimum cell numbers* in order to understand how atypia associations varied by the size of the urothelial cell clusters considered.

## Results

### Performance of Cell Border Detection Tool

Overall, the cell border detection tool was able to delineate the boundaries of negative and atypical urothelial cells with moderately high accuracy (*accuracy*=0.75; Table 3). Dense regions of indistinguishable cell borders were also detected with good accuracy (*IoU*=0.66; an *IoU* of 0.5 is considered good). However, there were instances where these dense regions were slightly overcalled (**Figure 2A,C,G,H**) and truncated (**Figure 1E, 2H**). Note in **Figure 2** instances where negative and atypical cells with distinct cell borders may have been folded into the dense regions (i.e., likely where regions were overcalled) and other instances where cells were nested in dense regions, suggesting the algorithm’s potential to conflate two distinct yet ambiguous entities. While this occurred in several instances, overall, this cross-contamination was not a significant issue (sensitivity=0.95, specificity=0.84; **Table 3**). Performance statistics for squamous and leukocytes were not reported as they were filtered out via the cluster postprocessing step.

**Table 3:**
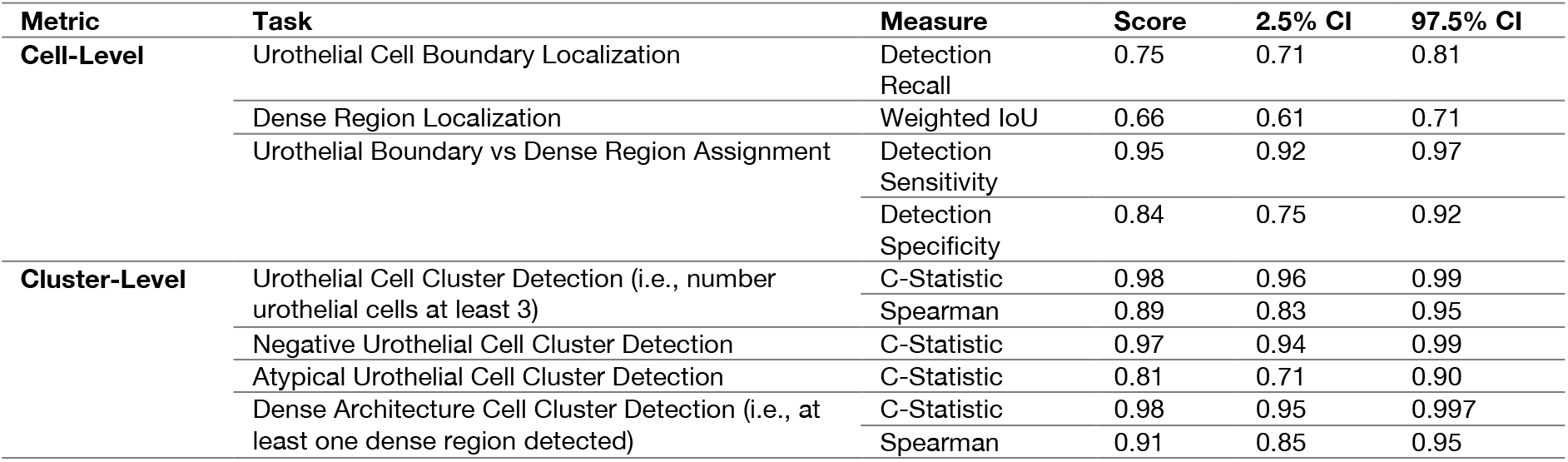
Performance metrics on internal test set of urothelial clusters.

**Figure 2:**
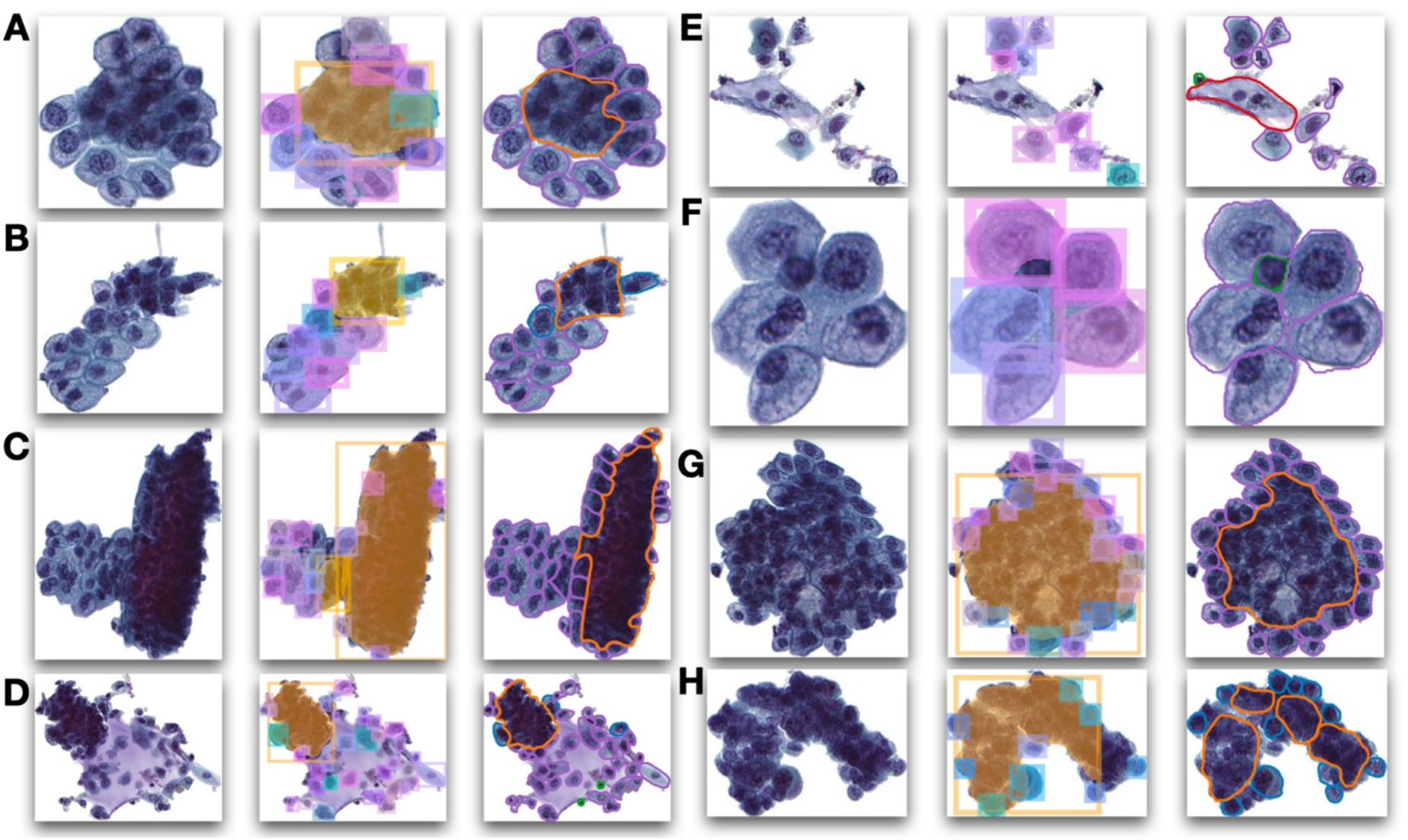
Examples of internal test set urothelial cell clusters: **A-H)** (left) original images; (middle) cell border detection model predictions with slight color jitter to denote distinct cell border predictions; squamous and inflammatory cells were removed from predictions during postprocessing as they were not the focus of this workflow; (right) pathologist annotations

In determining the relationship to overall UC atypia, only cluster-level aggregate measures (i.e., whether cluster harbors atypical cells or dense regions) are considered which do not require high fidelity pixelwise localization of the cell borders. Aggregate statistics of number of urothelial cells and dense regions were in excellent concordance with the ground truth annotations. The true and predicted number of urothelial cells in each cluster exhibited high correlation (*r=*0.89; **Table 3**), as did the dense region areas reported across the internal test set clusters (*r=*0.91; **Table 3**) (**Figure 3**). The cell border detection tool could accurately report on the presence of a urothelial cell cluster (i.e., number urothelial cells at least 3; C-Statistic=0.98), whether a cluster harbors atypical cells (C-Statistic=0.81) or whether a cluster contains a dense region (C-Statistic=0.98) (**Table 3**).

**Figure 3:**
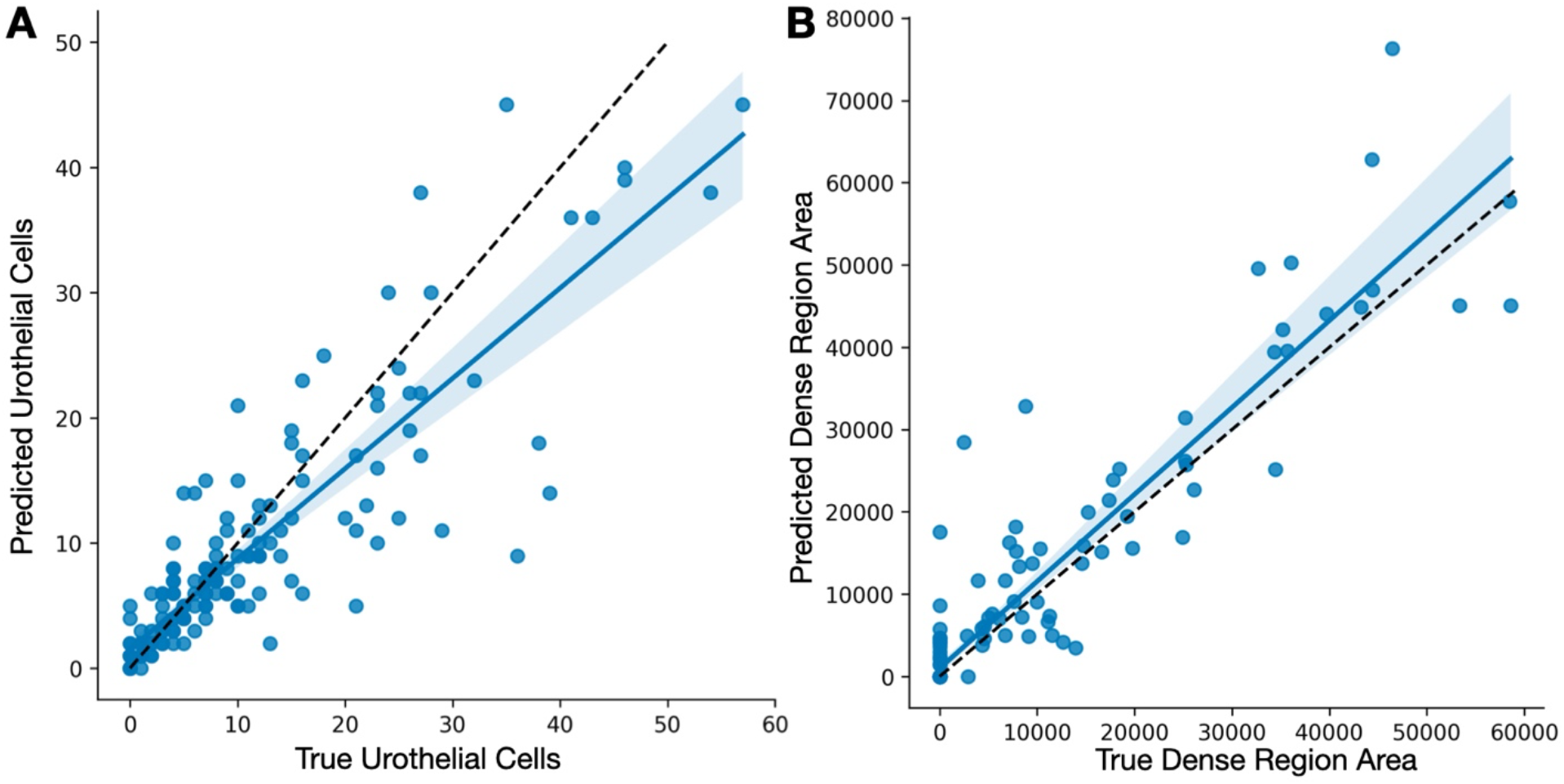
Scatterplots of aggregate true and predicted cluster statistics across internal test set clusters: **A)** counts of urothelial cells per cluster; **B)** area of dense regions within each cluster

### Associations with Overall UC Atypia

Presence and number of urothelial clusters was positively associated with specimen atypia, as were the number of clusters containing atypical cells or harboring dense regions (**Table 4; Figure 4**). However, these associations were modulated by the size of the urothelial cell clusters (**Figure 4A**). In general, the relationship between the number of urothelial/negative clusters and UC atypia strengthened based on the size of cluster considered. For instance, number of urothelial clusters containing only one urothelial cell or above (*minimum cell number*=1+; *IRR=1*.*53, 95%CI: [1*.*52-1*.*53, p<0*.*001;* **Table 4**) were less diagnostic for suspicious and positive as compared to number of clusters with 45 or more urothelial cells (*minimum cell number*=45+; *IRR=3*.*43, 95%CI: [2*.*42-4*.*82], p<0*.*001;* **Table 4**). Similarly, the number of atypical clusters containing only one urothelial cell or above (*minimum cell number*=1+; *OR=1*.*05, 95%CI: [1*.*04-1*.*06], p<0*.*001;* **Table 4**) were relatively nondiagnostic for suspicious and positive, whereas considering clusters with 45 or more urothelial cells were far more diagnostic (*minimum cell number*=45+; *OR=20*.*09, 95%CI: [3*.*56-137*.*02], p<0*.*001;* **Table 4**). The variance of the posterior credible interval of the effect estimates also changed based on the number of cells considered for a cluster (**Figure 4A**) as fewer clusters per specimen could be tabulated when more cells were required to call a cluster. There was a positive association between urothelial clusters that contained dense regions of overlapping cells and UC atypia (*minimum cell number*=8+; *OR=1*.*13, 95%CI: [1*.*07-1*.*19], p<0*.*001;* **Table 4**).

**Table 4:** Reported associations between urine atypia and number and type of urothelial clusters in specimen; depicted are the odds ratios for number of atypical urothelial clusters and number of dense urothelial clusters per specimen and incidence rate ratios for the total number of urothelial clusters per specimen; also included are the 95% credible intervals and “p-values”; results are reported by urothelial cluster size (*minimum cell number*)

| Type | Minimum Number of Urothelial Cells (i.e., Cluster Size) | OR/IRR | 2.5% CI | 97.5% CI | p |
| --- | --- | --- | --- | --- | --- |
| **Number Atypical Clusters** | 1+ | 1.05 | 1.04 | 1.06 | <0.001 |
|  | 2+ | 1.06 | 1.05 | 1.07 | <0.001 |
|  | 4+ | 1.14 | 1.12 | 1.17 | <0.001 |
|  | 8+ | 1.22 | 1.14 | 1.30 | <0.001 |
|  | 15+ | 1.86 | 1.51 | 2.27 | <0.001 |
|  | 21+ | 3.46 | 2.29 | 5.49 | <0.001 |
|  | 36+ | 10.66 | 3.59 | 34.99 | <0.001 |
|  | 46+ | 20.09 | 3.56 | 137.02 | <0.001 |
| **Number Dense Clusters** | 1+ | 1.13 | 1.13 | 1.14 | <0.001 |
|  | 2+ | 1.06 | 1.05 | 1.07 | <0.001 |
|  | 4+ | 0.90 | 0.88 | 0.92 | <0.001 |
|  | 8+ | 1.13 | 1.07 | 1.19 | <0.001 |
|  | 15+ | 1.13 | 0.94 | 1.36 | 0.19 |
|  | 21+ | 0.96 | 0.71 | 1.30 | 0.77 |
|  | 36+ | 0.88 | 0.45 | 1.69 | 0.692 |
|  | 46+ | 1.09 | 0.42 | 2.66 | 0.851 |
| **Number Total Clusters** | 1+ | 1.53 | 1.52 | 1.53 | <0.001 |
|  | 2+ | 1.61 | 1.60 | 1.62 | <0.001 |
|  | 4+ | 1.47 | 1.46 | 1.48 | <0.001 |
|  | 8+ | 1.26 | 1.24 | 1.28 | <0.001 |
|  | 15+ | 1.54 | 1.44 | 1.64 | <0.001 |
|  | 21+ | 1.65 | 1.49 | 1.83 | <0.001 |
|  | 36+ | 2.44 | 1.96 | 3.04 | <0.001 |
|  | 46+ | 3.43 | 2.42 | 4.82 | <0.001 |

**Figure 4:**
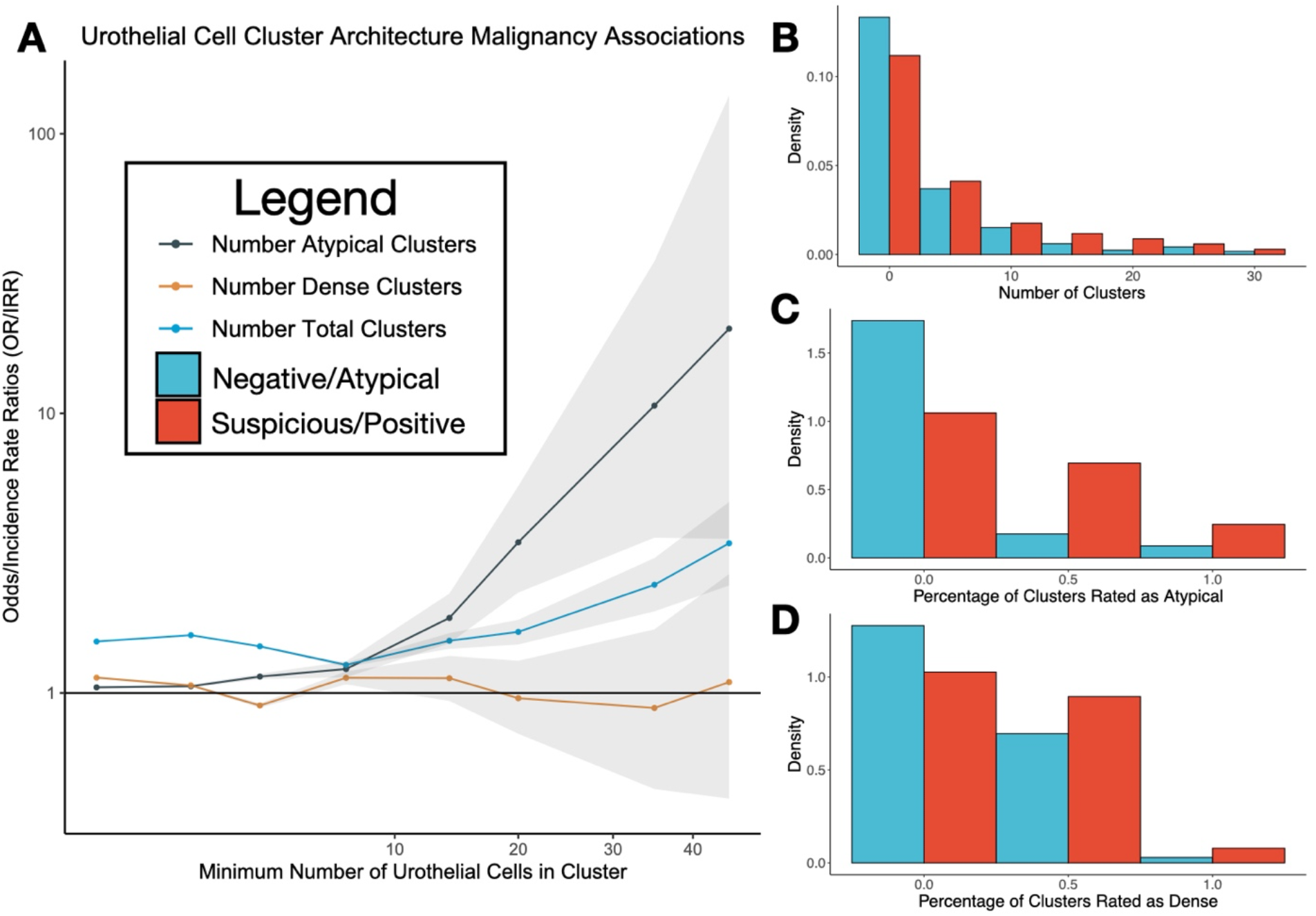
Graphical display of urine atypia associations with urothelial cell clusters: **A)** line plot depicting odds ratios/incidence rate ratios between specimen atypia and number of urothelial/atypical/dense clusters as a function of the minimum number of cells which form a urothelial cluster for recording number of clusters across specimen; horizontal line indicates no relationship (*OR/IRR*=1); grey regions denote 95% uncertainty/credible interval; **B)** grouped histogram normalized density plot of number of clusters per specimen versus whether specimen was suspicious/positive (blue) or negative/atypical (red), counts tabulated across specimen; plot was truncated to the right emphasize important relationships over outliers; demonstrates counts of urothelial cells were depleted for suspicious/positive specimen as compared to negative/atypical for lower number of cells and enriched at higher number of cells; *minimum cell number* was 20 for this example; **C)** similar plot for percentage of clusters within specimen that were rated as atypical; *minimum cell number* was 20 for this example; **D)** similar plot for percentage of clusters within specimen that contained a dense region; *minimum cell number* was 7 for this example; note that **C-D)** are unweighted by the total number of urothelial clusters in specimen

## Discussion

This work sought to better comprehend the association between cytological atypia for urothelial carcinoma and the number and type of urothelial cell clusters in voided urine specimens. While previous works have explored associations between presence, number and type of urothelial cell clusters and UC diagnoses, such studies were limited in nuanced exploration of these associations as they lacked the flexibility of a computer-based digital assessment. Meanwhile, existing computational methods for urine cytology do not clearly demarcate cellular boundaries nor do they explicitly define dense overlapping cellular architecture. The current work makes use of state-of-the-art deep learning methodologies to facilitate the incorporation of cluster atypia and architecture into the cytomorphological slide assessment by counting urothelial cells of atypical morphologies and enumerating distinct architectures (i.e., dense regions).

The study findings indicate that the cell border detection tool could accurately locate urothelial cells and dense overlapping pockets of urothelial cells within the specimen while disregarding squamous and inflammatory cells, which is a challenging task, even for a person. When evaluating cell clusters across the cohort, we recapitulated and expanded previous study findings on the importance of clusters for cytological assessment. The study findings presented in this paper confirmed the importance of assessing cytological atypia within clusters, though suggested that evaluating larger clusters may be more diagnostic ^31–33^. In concordance with previous findings, number of clusters within a specimen was found to be associated with specimen atypia, which may be reflective of the overall specimen cellularity. There exists ample literature documenting the effect of cellularity on specimen atypia ^2,4,34^. Meanwhile, this study hinted at potential relationships between the cell cluster architecture (i.e., presence of dense regions), which has been the subject of debate in existing literature, though these relationships were not as strong as the number of urothelial/atypical clusters.

While the cell border detection tool developed in this study identified associations between urothelial cell clusters and specimen atypia, it was never intended to be utilized as a diagnostic decision aid, but rather was intended to be incorporated into a more comprehensive algorithm as a preprocessing tool. While this study suggests the role and importance of clusters for cytology assessment, clearly the assessment of single cells is also equally if not more valuable for the final determination. Taken together, digital analysis of both individual cells and urothelial clusters may provide both efficient and reliable bladder cancer screening. However, there are several limitations to this study. For instance, this analysis was restricted to voided specimens prepared with ThinPrep®, and results may differ depending on the specimen type and preparation method. Other confounders and/or effect modifiers (e.g., age, sex, previous history of hematuria, stones, bladder cancer, biopsy results) may influence the associations explored in this study and warrant future exploration. As there were significantly fewer suspicious and positive diagnoses than atypical and negative, this could point to potential selection bias, especially since the number of atypical and dense clusters were only compared between groups for cases where the number of total urothelial clusters was greater than zero. Such cases were still included for findings based on number of clusters, although the number of clusters per specimen decreases as the minimum number of cells to call clusters increases (i.e., zero-inflated outcome). Findings were guided by evaluation using TPS, were done in research setting and did not incorporate information from the medical history and cytotechnologist prescreening, all of which may limit the external applicability of the study findings. Additionally, different cytopathologists may evaluate clusters differently, which presents a future area of exploration. There also exist other cell types which may have evaded assessment (e.g., seminal vesicle cells, glandular cells, etc.). Folding in atypical cells into dense clusters may also have impacted the final assessment. In some cases, pathologist annotations of the cell boundaries within clusters were coarse, which may have distorted reporting of the detection tool accuracy. Collecting additional training data and exploring new methods and other augmentation techniques present opportunities to improve the detection model.

Over the past decade, urine cytology assessment criteria have become increasingly quantitative to resolve interobserver variability in atypical and suspicious cytology assignments. Reporting reliability and promptness, as well as disease management options may be further improved through assessment of all cells within the specimen, though manual assessment on this scale is currently intractable and unfeasible given the caseload. The preprocessing cell border identification/localization tool could serve as an important upstream step for a diagnostic decision support aid, which could operate on candidate urothelial cells extracted within clusters to provide reliable atypia estimates (e.g., precisely estimate NC ratio for confirmed urothelial cells within cluster).

## Conclusion

Building a comprehensive understanding of the relevance of urothelial cell cluster atypia for cytological bladder cancer screening is a challenging task as it requires the precise localization of cell borders within complex cellular mixtures of varying overlap. For emerging digital diagnostic aids, assessment of clusters remains an ambiguous and *ad hoc* accompaniment to single cell analysis. This study sought to develop a deep learning-based preprocessing tool for separating cell borders and where appropriate, registering the presence and location of dense, highly cellular architectures. While the current study pointed to associations between cluster atypia and urothelial carcinoma, we plan to incorporate the cell border detection tool into a digital workflow for rapid bladder cancer screening to investigate the degree to which such a tool can augment clinical decision making.

