## Appendix for "Uncovering Additional Predictors of Urothelial Carcinoma from Voided Urothelial Cell Clusters Through a Deep Learning Based Image Preprocessing Technique"

### Supplementary Methods

#### Modeling Number and Type of Urothelial Clusters

Number of urothelial clusters was modeled as a Poisson distributed outcome as the number of clusters per specimen is at least 0 and unbounded, whereas number of atypical and dense clusters were modeled as a Binomial distributed outcome as the number of atypical/dense clusters is bounded by the total number of clusters per specimen, making the target of inference the proportion of clusters that are atypical/dense:

$$\text{number clusters}_{\mathrm{i}} \sim \text{Poisson}(\lambda_{\mathrm{i}})$$
$$log(\lambda_i) = \beta_0 + \beta_1 \mathbb{1}_{x_i \in \{sus.pos\}} + \theta_{pathologist[i]} + \theta_{patient[i]}$$

$$\text{number atypical/dense clusters}_i \sim Binomial(\text{number clusters}_i, p_i)$$
$$\text{logit}(p_i) = \beta_0 + \beta_1 \mathbb{1}_{x_i \in \{sus.pos\}} + \theta_{pathologist[i]} + \theta_{patient[i]}$$

$$\theta_{pathologist[i]} \sim N(0, \tau_1{}^2)$$
$$\theta_{patient[i]} \sim N(0, \tau_2{}^2)$$
$$\beta \sim N(0,4)$$

Clustering by cytopathologist and nesting within patient across multiple longitudinal assessments was accounted for through the addition of random intercepts. From these models, we were able to report incidence rate ratios (*IRR*) and odds ratios (*OR*), calculated by $\text{IRR}|\text{OR} = \exp(\beta_1)$, to determine whether suspicious or positive ratings were associated with greater or lesser number of clusters as compared to atypical and negative ratings. High density posterior credible intervals (CI) (analogous to the confidence interval) for these ratios were given through sampling the posterior distribution 4000 times across four Markov Chains.
